# Behavioural traits and feeding ecology of Mediterranean lionfish and native species naiveté to lionfish predation

**DOI:** 10.1101/2020.01.22.915322

**Authors:** Daniele D’Agostino, Carlos Jimenez, Tom Reader, Louis Hadjioannou, Stephanie Heyworth, Marilena Aplikioti, Marina Argyrou, David A. Feary

## Abstract

The detrimental effects of invasion by Indo-Pacific lionfish (*Pterois volitans/miles*) on western Atlantic fishes have spurred concerns for Mediterranean fish biodiversity, where a Lessepsian invasion of lionfish has recently begun. In order to assess the potential impact on biodiversity, we examine key behavioural and ecological traits of lionfish, and the resident fish community in the Mediterranean, that may contribute to lionfish invasion success. We focus on Cyprus, where lionfish populations were first sighted in 2012 and have now established abundant and stable populations. Using field observations, we examine lionfish predatory behaviour and feeding ecology, and resident fish species naiveté to hunting lionfish. Our findings suggest that lionfish in the Mediterranean are crepuscular generalist predators, with prey targeted dominated by small-bodied benthic or bentho-pelagic associated species. Such prey are more likely to be native than introduced (Lessepsian) fishes, with native prey fishes showing greater naiveté towards lionfish than Lessepsian prey species. Notably, one of the Mediterranean’s key ecological fish species (the native damselfish *Chromis chromis*), showed the highest level of naiveté and was the most heavily targeted prey. Overall, lionfish in the Mediterranean show similar predatory behaviour and ecology to their western Atlantic counterparts. Although the Mediterranean invasion is still relatively recent, it may result in a similar disruption to reef fish biomass to that recorded in the Atlantic, with impact to the structure and biodiversity of reef fish communities and the services they provide.

## 1. INTRODUCTION

Human activities, such as land-cover change, chemical release, overharvesting, climate change and species transport/invasion, are having a profound impact on biodiversity worldwide (Crutzen 2006, Dirzo et al. 2014, McGill et al. 2015, Gordon et al. 2018). Invasive species, which often constitute new functional components in the recipient community, can generate ecological impacts that propagate along the food-web triggering trophic cascades (Moyle & Light 1996, Strayer 2010) and result in a generalised decrease in the abundance and diversity of native communities (Gallardo et al. 2016). Therefore, a central focus of invasion biology research has been to examine the factors which increase our ability to predict which species may become invasive (Kolar & Lodge 2001, Hayes & Barry 2008). This work has shown that there are a range of life-history, ecological and behavioural traits that are associated with successful invasion, and that can be used to determine the subsequent consequences of an invasion on the invaded ecosystem (Chapple et al. 2012).

The invasion process involves a series of sequential stages (transport, introduction, establishment and spread) that a population has to go through to become a successful invader (Blackburn et al. 2011, Chapple et al. 2012). Each stage presents varied abiotic (i.e. associated with different geography and environment) and biotic barriers (i.e. dispersal, reproduction, feeding, competition) that individuals within an invading population must overcome to succeed (Blackburn et al. 2011); only individuals with certain life-history and behavioural traits may then move forward between steps in the invasion process (Chapple et al. 2012). As a result of such ‘filtering’ of individuals, the invading population is not a random subset of the native natal population (Felden et al. 2018), while traits ‘selected’ for in individuals during one stage of the invading process may being maladaptive and lead to invasion failure in following stages (Chapple et al. 2012, Felden et al. 2018). Thus, the introduction or establishment of an invasive species in a novel region will not necessary lead to the species becoming invasive and to similar impacts on local biodiversity as observed elsewhere.

Substantial negative impacts on native fish biodiversity have been well documented following the invasion of lionfish *Pterois volitans/miles* (hereafter ‘Atlantic lionfish’) in the western Atlantic ocean (Albins & Hixon 2013, Côté & Smith 2018). Such biodiversity loss may mirror that for native biodiversity in the Mediterranean (Galanidi et al. 2018), where a Lessepsian invasion (i.e. from the Suez Canal, Egypt) by lionfish (*P. miles*, [Bennett, 1828], hereafter ‘Mediterranean lionfish’) has recently begun (Bariche et al. 2013, Jimenez et al. 2016, Kletou et al. 2016, Azzurro et al. 2017). Research on Mediterranean lionfish is still in its infancy, and is still not clear whether Mediterranean lionfish are set for a pan-Mediterranean invasion or whether they will be limited to the Levantine side of the basin (Johnston & Purkis 2014, Azzurro et al. 2014), where the environmental affinity to the Red Sea is higher (i.e. warmer water temperature). Nevertheless, the extensive work on the ecological, behavioural and life-history traits contributing to invasive success of lionfish in the western Atlantic provides a basis for understanding the traits potentially facilitating the Mediterranean invasion (reviewed by Côté & Smith 2018). Such understanding may allow us to predict the impact on local and regional biodiversity of the Mediterranean invasion, and the management decisions needed to effectively mitigate long term changes in community structure (Morris & Whitefield 2009).

Mediterranean lionfish share important life-history and trophic traits with Atlantic lionfish, which may enhance the likelihood of invasion success within the Mediterranean (Kleitou et al. 2019). High growth rates (up to 20 cm total length [TL] in the first year), large body size (up to 37 cm TL at four years) and early maturation (sexual maturity in less than a year) in Mediterranean lionfish (Kleitou et al. 2019), can be expected to translate into rapid rates of population increase (Morris & Whitefield 2009, Edwards et al. 2014, Côté & Smith 2018), which is often an essential prerequisite to outcompete native competitors for spaces and resources during the establishment phase of the invasive process (Holway & Suarez 1999, Chapple et al. 2012). Moreover, early studies on Mediterranean lionfish stomach contents report that they feed on a wide range of crustacean and fish species (Kleitou et al. 2019, Zannaki et al. 2019), suggesting they are generalist predators and that they may be well equipped to deal with environmental stochasticity in the invaded region (García-Berthou 2007, Peake et al. 2018). However, evidence of dietary specialization on species with particular morphological (i.e. small, elongated body) and behavioural traits (i.e. benthic habitat use, solitary) have been reported for the Atlantic lionfish (Green & Côté 2014, Chappell & Smith 2016), implying stronger predatory pressure on certain species, increasing the risk of localised species extirpation and loss of entire ecological and functional roles (Peake et al. 2018). Lastly, although there is no information on native prey naiveté to Mediterranean lionfish, results from the Atlantic would suggest that naiveté will be an important precursor to Mediterranean lionfish population success (Anton et al. 2016, Benkwitt 2017, Haines and Côté 2019). In detail, the ‘naïve prey’ hypothesis postulates that native prey suffer heavy predation by a novel predator due to the lack of recent co-evolutionary history and effective antipredator behaviour (Cox & Lima 2006, Sih et al. 2010, Paolucci et al. 2013). Such prey naiveté may then allow the invasive predator a competitive advantage compared to native counterparts (Anton et al. 2016).

To predict how the structure and biodiversity of Mediterranean fish communities will be impacted, the objective of this study was to document and assess patterns in activity and feeding of Mediterranean lionfish, as well as investigate the potential for native (compared to non-native) prey naiveté. We focus on Cyprus, which is at the forefront of the Mediterranean invasion and holds an abundant and stable population of lionfish in shallow and deep waters (Jimenez et al. 2016, 2019a, Kletou et al. 2016). First (i), we quantified the behaviour of Mediterranean lionfish at three time points (i.e. sunrise, noon and sunset) to test the hypothesis of a crepuscular pattern in hunting activity (i.e. sunrise and sunset) as this has been shown in the Atlantic invasion (Cure et al. 2012, Benkwitt 2016) and invoked as one of the behavioural traits facilitating their invasive success (i.e. increased hunting success and broader array of preys available) (Hobson 1973, Potts 1990, Green et al. 2011).

Secondly (ii), we characterised the feeding ecology of Mediterranean lionfish, quantifying the diversity of species preyed upon, and the number of times each species was targeted. We relate this feeding behaviour to fine-scale and broad-scale prey abundance. We predicted that lionfish would broadly be generalist predators by feeding on a diverse array of prey (Peake et al. 2018), but would also show some level of dietary specialization towards species with certain behavioural and morphological traits (Green & Côté 2014, Chappell & Smith 2016). Lastly (iii), we investigated the levels of behavioural naiveté towards lionfish in native and Lessepsian prey species, predicting that native prey would be more naïve towards hunting lionfish (Anton et al. 2016).

## 2. MATERIAL & METHODS

### 2.1. Site selection and methodology for recording and quantifying Mediterranean lionfish behaviour

To develop a dataset of Mediterranean lionfish behaviour, in September 2018 we videoed Mediterranean lionfish behaviour at sunrise (between 7:30-8:30), noon (13:30-14:30) and 2 sunset (17:00-18:00) across six rocky reefs (mean area 0.056 km^2^ ± 0.004 SE) off the eastern coast of Cyprus (Protaras; 35° 0’5.65”N, 34° 4’11.88”E) (Fig. 1 for details). A total of 80 individual Mediterranean lionfish were recorded across sites and time of the day. All sites were dominated by rocky reef and ranged in depth between 3.8 and 12.3 m (mean depth 7.5 m ± 0.2 SE). Sites were separated by at least 600 m of sand and deeper waters (up to 30 m). Sites and time of observations were randomised. Within each site where Mediterranean lionfish were identified, care was taken to not video the same individual twice by swimming unidirectionally along each site during the dive, and videoing fish as encountered (Green et al. 2011). Each site was dived only once within a day (∼1 hour dive), and if we returned to the same site on a different day, care was taken to observe fish at different time periods from the previous visit (i.e. sunrise, noon or sunset) and in different areas of the site (Côté & Maljkovic 2010, Cure et al. 2012).

**Fig. 1.**
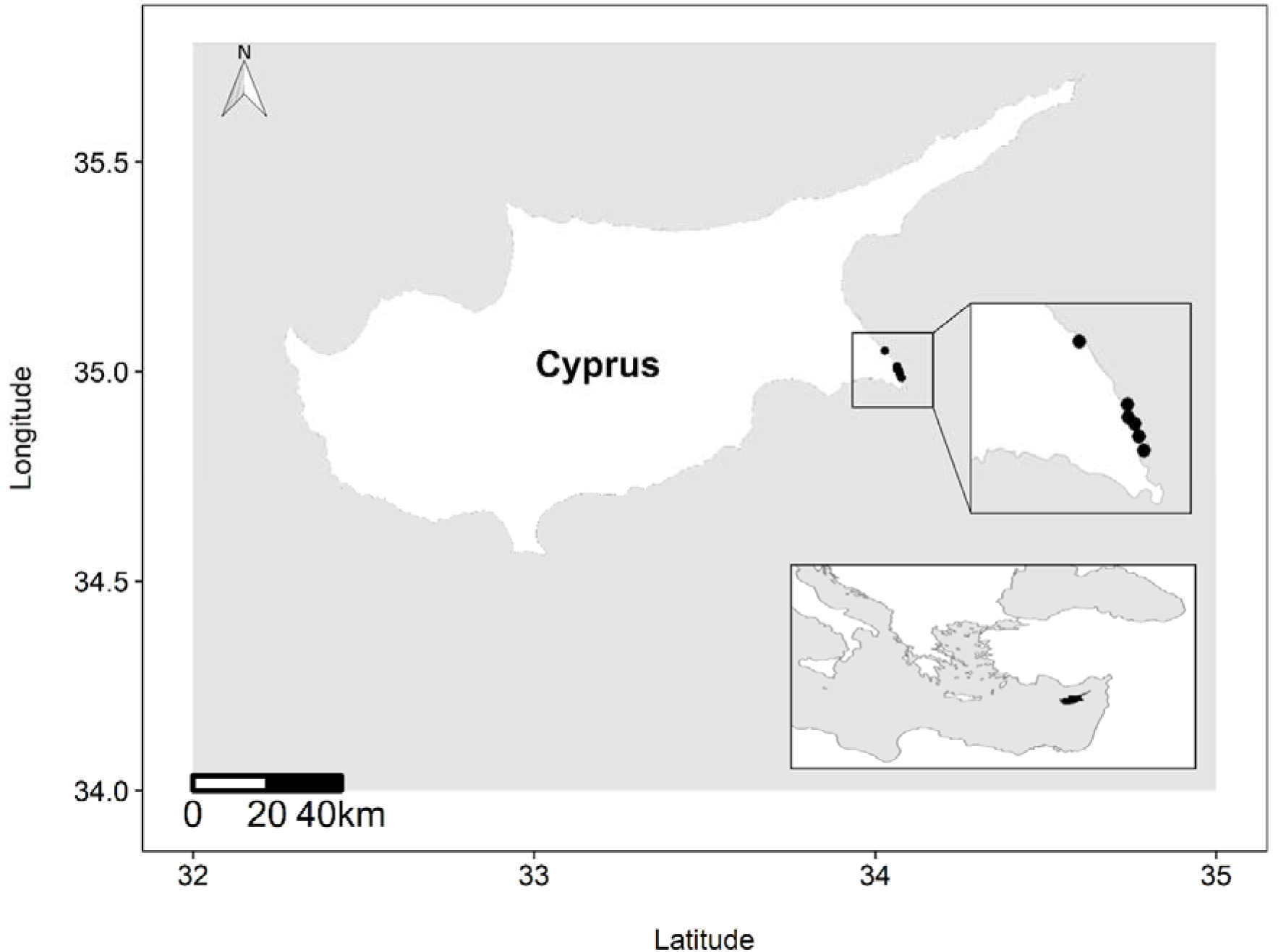
Study area and sampling locations of the western part of Cyprus.

Once identified, Mediterranean lionfish were allowed to acclimate for 3 min to the diver, and were then filmed for 5 min, with a minimum distance of 3 m between diver and lionfish (D’Agostino et al. 2019). During the 3 min acclimation period the diver estimated Mediterranean lionfish body size (small [≤ 10 cm TL], medium [11 – 20], large [≥ 21]), group size (number of lionfish within 1 m radius from focal lionfish), water depth (m) and temperature (nearest 0.1 °C), measured using the dive computer RATIO ix3M, visibility (low < 10 m, medium 10 – 20 m, high > 20 m) and current strength (low: diver barely kicking to maintain position, medium: periodic kicking required by diver to maintain position, high: constant kicking by diver required to maintain position); cloud cover was recorded at the beginning of each dive as either clear: 0 – 25%, partly cloudy: 25 – 75% or overcast: > 75% (Côté and Maljkovic 2010; Cure et al. 2012). Underwater data collection and video recording were carried out by three experienced divers (DD, SH and LH); pilot data (not included in the study) where all divers observed the same individuals, were collected on the first day and used to calibrate data collections methodology between observers. Mediterranean lionfish size estimation error was < 10 %.

To quantify activity budget of each Mediterranean lionfish, videoed behaviour was classified into four categories, with each category of behaviour scored every 10 seconds across the 5 min video (Green et al. 2011). Mediterranean lionfish were considered ‘resting’ when motionless on the substratum, with dorsal spines held flat along the dorsal midline and pectoral fins closed, ‘hunting’ when approaching/chasing potential prey, with head and flared pectoral fins directed at prey and dorsal spines undulating and erected, ‘hovering’ when nearly motionless above the substratum, and ‘transiting’ when swimming from one part of the benthos to another (see supplementary materials Video S1-4 for examples of lionfish resting and hunting behaviour). In addition, to quantify movement of each Mediterranean lionfish ‘distance moved’ was quantified (in cm) by estimating the total distance moved (to the nearest 5 cm) every 30 sec interval across each 5 min video, using the lionfish individual size as reference against the background reef. Any videos in which fish displayed an adverse reaction to the diver’s presence (i.e. staring at divers or assuming defensive posture, < 5 % of individuals) were excluded from analysis.

### 2.2. Quantification of Mediterranean lionfish feeding ecology and fish community structure

To quantify Mediterranean lionfish feeding ecology, across each 5 min video we counted the number of times individuals (hereafter ‘prey species’) were targeted (‘times targeted’) across all predation attempts (defined as the action of a lionfish pursuing a single individual). Prey species trophic guild and origin (i.e. native or Lessepsian) were extracted from FishBase (Froese & Pauly 2000).

To characterise the composition of the community with which Mediterranean lionfish associate, either because of shared habitat preferences, or perhaps because of predator-prey relationships, the abundance of all prey species within 1 m radius of focal lionfish was estimated within each video (hereafter ‘fine-scale prey abundance’). To characterise the fish community in the broader area to which Mediterranean lionfish could potentially associate or prey upon, ‘broad-scale prey abundance’ was estimated as the sum of all Mediterranean lionfish prey estimated from underwater visual census surveys conducted on SCUBA during six sampling expeditions (winter, spring, summer and autumn 2017; winter and summer 2018). Such surveys were undertaken at five reef sites (between Protaras and Cape Greco, Fig. S1) in the same area as where behavioural observations were quantified. Sites were located at depths ranging between 3.5 and 20 m (average 9.3 m). At each site three replicate transects of 25 x 5 m were surveyed and all fish identified to species, enumerated and their size (cm TL) estimated. Only prey fish ≤ 5 cm TL and from species that had been previously observed to have been targeted by lionfish during the behavioural observations, were selected to estimate fine- and broad-scale prey abundance.

### 2.3. Quantification of prey fish naiveté to Mediterranean lionfish predation

To examine and compare naiveté between native Mediterranean and Lessepsian prey species, the ‘closest approach distance’ (i.e. the distance [cm] at which small prey fish [i.e. ≤ 5 cm TL] stopped approaching or turned away from lionfish) was quantified during each 5 min video (Anton et al. 2016). Distance between prey and Mediterranean lionfish was visually estimated by using focal Mediterranean lionfish size (which had been previously estimated in situ) as a reference. If the same prey individual approached a Mediterranean lionfish multiple times during the 5 min observation period, only the closest approach was enumerated. The distance of 60 cm was used as minimum starting distance for observation of closest approach distance, as pilot observations showed that prey species predominantly respond to the presence of hunting lionfish within this distance. In order to control for predator behaviour, closest approach distance measures were only calculated from videos where Mediterranean lionfish were identified as hunting (see below) (n = 31).

Mediterranean lionfish behaviour, feeding ecology and native prey naiveté data were initially extracted from 15% of videos and independently quantified by two observers. As results differed by < 5% between observers, all remaining quantification of data from videos analyses were carried out by one observer (DD).

### 2.4. Statistical analysis

#### 2.4.1 Behavioural patterns of Mediterranean lionfish

As there was no significant difference in visibility (always greater than 20 m), currents (always low), cloud cover (always clear) and depth (Kruskal-Wallis test, H = 3.3323, df = 2, p = 0.189) between sampling periods (sunrise, noon, sunset), these environmental variables were not included in further statistical models. Similarly, water temperature was excluded from further analysis because its variation between sampling periods, although statistically significant (H = 15.328, df = 2, p < 0.001), was very limited in magnitude (27.0 to 28.2°C), reflecting normal diel temperature variation during the observational period.

Behavioural analyses only focused on resting and hunting, which represented > 80 % of the behaviours enacted across all videos, as well as total distance moved. To understand the relationship between the response variables of resting, hunting and total distance moved to the independent factors of sampling time (sunrise, noon and sunset), prey availability, Mediterranean lionfish body size (small, medium and large) and group size (one, two or three individuals), separate generalised linear mixed models (GLMMs) were used. A GLMM with binomial error distribution and logit link function was used to model the probability of resting and hunting behaviour being observed in a 5 min video. We took this approach because data for proportion of time spent resting and hunting were bimodal, with a high proportion of ones and zeros, and data transformation did not result in either normality or homoscedasticity within either data sets. An individual was considered as ‘resting’ if > 50 % of the proportion of time during the 5 min period was spent resting, and ‘hunting’ if > 50 % of the 5 min period was spent hunting (Cure et al. 2012). To model total distance moved a Gaussian error structure was used. Total distance moved data were log (X + 1) transformed prior to testing to satisfy test assumptions. In both GLMMs, we fitted random intercepts for the factors site (six levels) and date of sampling (five levels), with backwards model selection then followed using likelihood ratio tests to examine the significance of each term removed from each model.

#### 2.4.2 Mediterranean lionfish feeding ecology and prey association

To determine whether Mediterranean lionfish were associated with, and predated upon, prey in relation to their abundance in the wider community, two types of test were conducted. First, we used Pearson’s correlation tests to look for a relationship between the likelihood of a prey species being targeted, its local (fine-scale) abundance, and its broad-scale abundance in the area. A positive correlation would imply that prey selection is influenced by availability. Second, we used chi-squared tests to determine whether there were significant differences in the relative abundance of prey species targeted by Mediterranean lionfish, and their relative fine- and broad-scale abundance. A significant difference would imply that Mediterranean lionfish are selective about the prey that they attack. In order to meet chi-squared test assumptions, *Thalassoma pavo*, *Coris julis* and *Symphodus* spp. individuals were grouped together by family as ‘Labridae’; *Gobius vittatus* (Gobiidae), *Scorpaena notata* (Scorpaenidae) and *Apogon imberbis* (Apogonidae) were grouped as ‘rare species’; while *Siganus rivulatus* (Siganidae) and *Sparisoma cretense* (Scaridae) were grouped together because they had relatively high abundance, but were scarce in times targeted (Table S1).

**Table 1.**
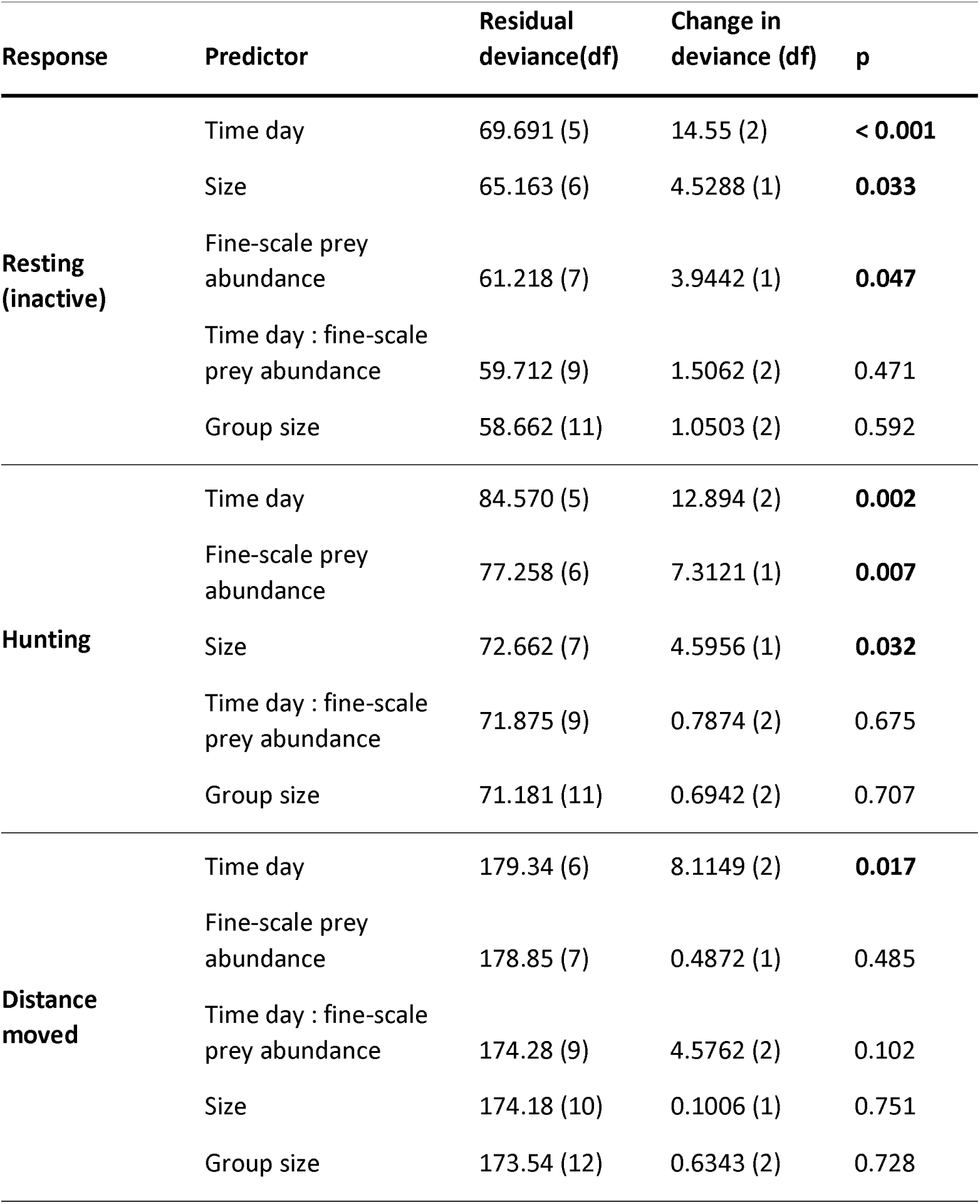
Result of GLMM analysis of the effects of biotic/abiotic factors on Mediterranean lionfish’s behaviour. Binomial error distribution with logit link function for resting and hunting behaviour; Gaussian error structure for distance moved. Significant p values highlighted in bold.

| Response | Predictor | Residual deviance(df) | Change in deviance (df) | p |
| --- | --- | --- | --- | --- |
| **Resting (inactive)** | Time day | 69.691 (5) | 14.55 (2) | **< 0.001** |
|  | Size | 65.163 (6) | 4.5288 (1) | **0.033** |
|  | Fine-scale prey abundance | 61.218 (7) | 3.9442 (1) | **0.047** |
|  | Time day : fine-scale prey abundance | 59.712 (9) | 1.5062 (2) | 0.471 |
|  | Group size | 58.662 (11) | 1.0503 (2) | 0.592 |
| **Hunting** | Time day | 84.570 (5) | 12.894 (2) | **0.002** |
|  | Fine-scale prey abundance | 77.258 (6) | 7.3121 (1) | **0.007** |
|  | Size | 72.662 (7) | 4.5956 (1) | **0.032** |
|  | Time day : fine-scale prey abundance | 71.875 (9) | 0.7874 (2) | 0.675 |
|  | Group size | 71.181 (11) | 0.6942 (2) | 0.707 |
| **Distance moved** | Time day | 179.34 (6) | 8.1149 (2) | **0.017** |
|  | Fine-scale prey abundance | 178.85 (7) | 0.4872 (1) | 0.485 |
|  | Time day : fine-scale prey abundance | 174.28 (9) | 4.5762 (2) | 0.102 |
|  | Size | 174.18 (10) | 0.1006 (1) | 0.751 |
|  | Group size | 173.54 (12) | 0.6343 (2) | 0.728 |

#### 2.4.3. Native versus invasive prey naiveté to Mediterranean lionfish predation

To determine whether there was a significant difference in closest approach distance between native and invasive prey species and Mediterranean lionfish, and whether prey species would approach lionfish at different distances in regards of their size, a GLMM with Poisson error distribution and log link function was used. The model had closest approach distance as the response variable and the interaction between origin (native vs invasive) and lionfish size (small, medium, large) as fixed effects, with lionfish ID as random intercept (as multiple observations were conducted on each lionfish). All data were analysed in R (R core development team 3.5.1, 2019) using the nlme package (Pinheiro et al. 2015) for mixed-effects models, and statistical significance was determined at p-values < 0.05.

## 3. RESULTS

### 3.1. Behavioural patterns of Mediterranean lionfish

We quantified patterns in activity and feeding of 76 Mediterranean lionfish (sunrise n = 27, noon n = 25 and sunset n = 24) ranging in size from 7 to 35 cm TL (mean ± SE: 19 ± 0.8 cm) during a total of 380 min of video. Mediterranean lionfish were observed alone or in groups of up to three individuals. Time of the day had a significant effect on resting, hunting and distance moved (Table 1), with Mediterranean lionfish spending the highest proportion of time hunting, as well as moving longer distances at sunrise and sunset (Fig. 2, Fig. 3a). At noon Mediterranean lionfish spent the majority of time resting (supplementary materials Video 1). Lastly, fine-scale prey abundance and Mediterranean lionfish individual size had a significant effect on resting and hunting behaviour, with hunting behaviour more likely when prey availability was high and lionfish size was small (Table 1). Fine-scale prey abundance ranged between 0 to 41 individual prey fishes per Mediterranean lionfish, with highest abundances apparent at sunset (Fig. 3 b).

**Fig. 2.**
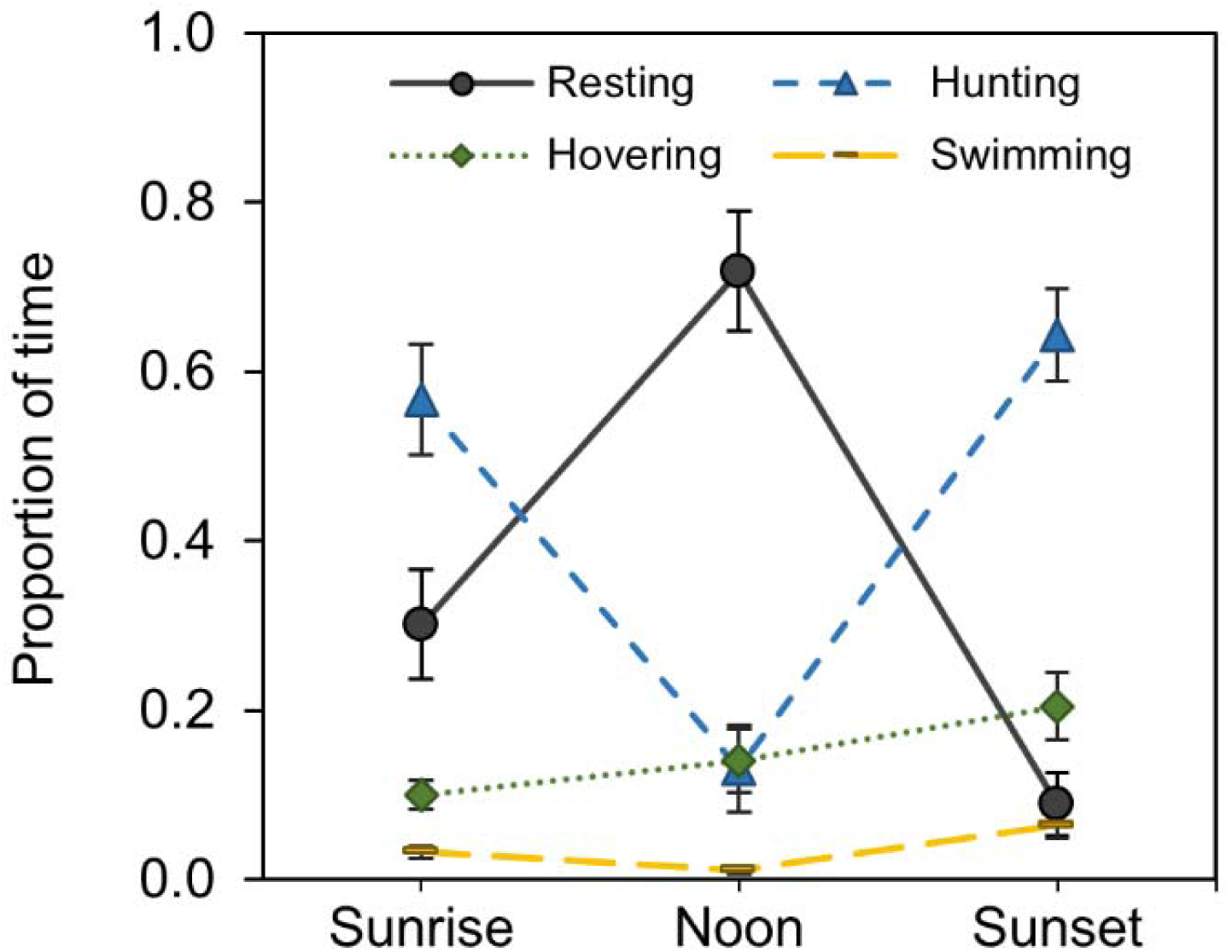
Time budget of invasive Mediterranean lionfish on Cypriot rocky reefs across time of day for resting, hunting, hovering and swimming. Shown are mean (± SE) proportions of time spent in each behaviour.

**Fig. 3.**
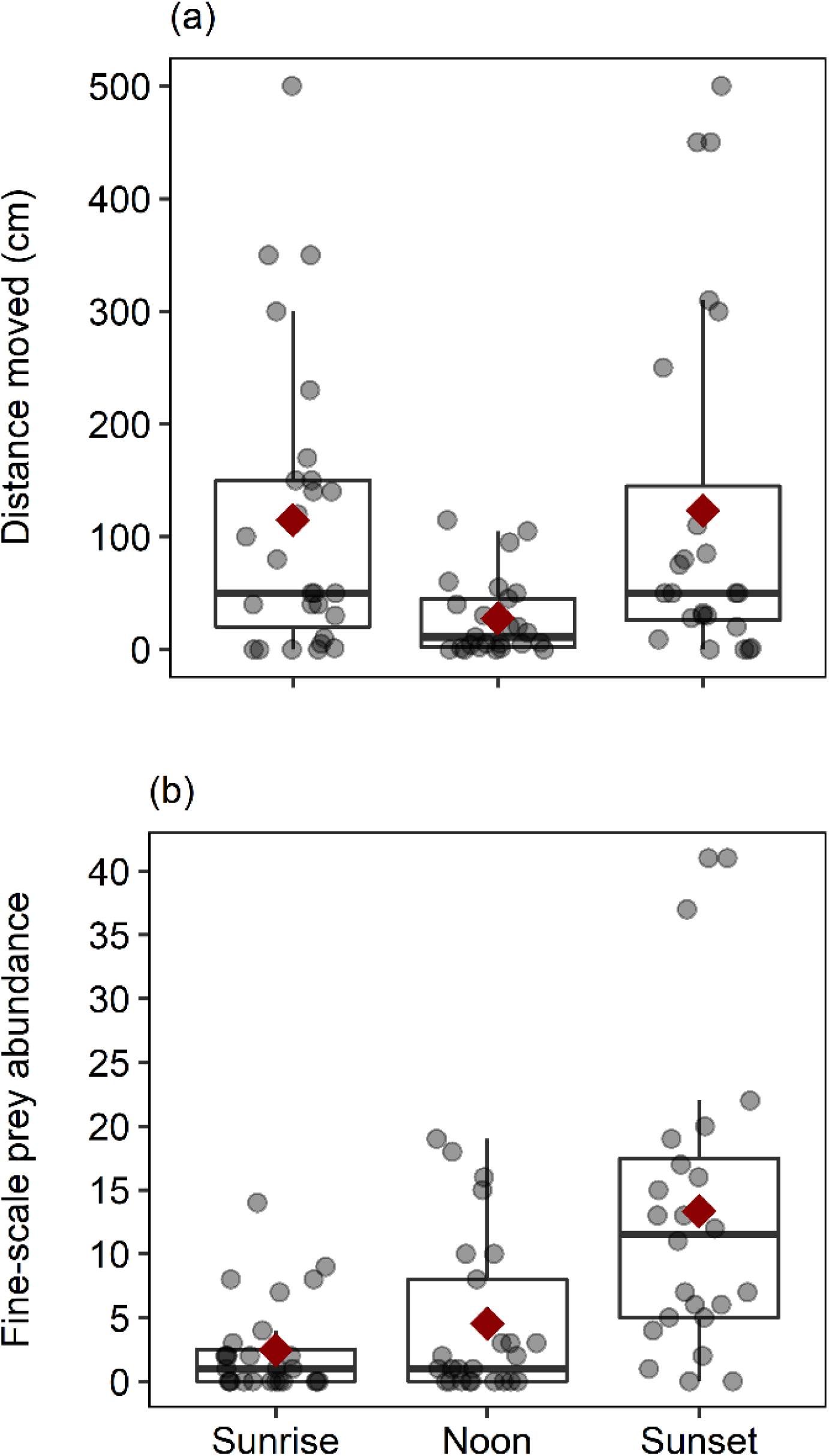
(a) Changes in Mediterranean lionfish distance moved among time of day and (b) changes in fine-scale prey abundance at different time of the day. Thick horizontal line shows the median, boxes show inter-quartile (IQR). Whiskers indicate the range of data, dots show individual data point, and red diamond show the mean.

### 3.2. Mediterranean lionfish feeding ecology and prey association

Mediterranean lionfish targeted 10 prey species from seven families, encompassing only grazers and meso-predators (Table 2). Prey targeted were predominantly small-bodied benthic or bentho-pelagic associated species; predatory activity was dominated by predation attempts on *Chromis chromis* (Pomacentridae) and juvenile *T. pavo* (Labridae) (supplementary materials Video 2 and 3, respectively).

**Table 2.** Potential prey fish species, the frequency with which they were targeted by Mediterranean lionfish, and their relative abundance in the immediate vicinity of Mediterranean lionfish (fine-scale abundance) and in the broader area (broad-scale abundance). Both absolute (total number of individuals

| Targeted species | Family | Trophic level (± SE) | Origin | Life status | Times targeted | Fine-scale prey abundance | Broad-scale prey abundance | Time targeted index (%) | Fine-scale prey relative abundance (%) | Broad-scale prey relative abundance (%) |
| --- | --- | --- | --- | --- | --- | --- | --- | --- | --- | --- |
| *Chromis chromis* † | Pomacentridae | 3.8 (0.4) | Native | Adult | 42 | 331 | 6162 | 48.28 | 64.77 | 63.99 |
| *Thalassoma pavo* | Labridae | 3.5 (0.47) | Native | Juvenile | 22 | 64 | 193 | 25.29 | 12.52 | 2 |
| *Parupeneus forsskali* | Mullidae | 3.5 (0.3) | Invasive | Juvenile | 8 | 50 | 336 | 9.2 | 9.78 | 3.49 |
| *Apogon imberbis* | Apogonidae | 3.4 (0.61) | Native | Adult | 4 | 6 | 5 | 4.6 | 1.17 | 0.05 |
| *Coris julis* | Labridae | 3.4 (0.1) | Native | Juvenile | 4 | 13 | 330 | 4.6 | 2.54 | 3.43 |
| *Sparisoma cretense* | Scaridae | 2.9 (0.24) | Native | Juvenile | 2 | 8 | 619 | 2.3 | 1.57 | 6.43 |
| *Gobius vittatus* † | Gobiidae | 2.9 (0.27) | Native | Adult | 1 | 1 | 2 | 1.15 | 0.20 | 0.02 |
| *Siganus rivulatus* | Siganidae | 2.0 (0.0) | Invasive | Juvenile | 1 | 34 | 1966 | 1.15 | 6.65 | 20.42 |
| *Symphodus sp.* | Labridae | - | - | - | 1 | 2 | 5 | 1.15 | 0.39 | 0.05 |
| *Symphodus rostratus* | Labridae | 3.5 (0.0) | Native | Juvenile | 1 | 1 | 2 | 1.15 | 0.2 | 0.02 |
| *Scorpaena notata* | Scorpaenidae | 3.7 (0.2) | Native | Juvenile | 1 | 1 | 9 | 1.15 | 0.2 | 0.09 |

sighted across all samples) and relative (proportion of all individuals sighted) measures of abundance are given.

Although important, community prey abundance did not completely predict Mediterranean lionfish species-specific targeting rate. Despite the significant positive correlation between the number of times a species was targeted and its fine-scale abundance, and between fine-scale and broad-scale prey abundance, the likelihood of a species being predated upon was not always proportionate to its abundance, either in the immediate vicinity of the lionfish, or in the broader area (Table 3). For example, while *C. chromis* broad-scale abundance (64%) reflected its fine-scale abundance (64.8%) and the number of times it was targeted (48.3%), we observed contrasting trends for *S. rivulatus/S. cretense* and Labridae. In detail, despite *S. rivulatus/S. cretense* being the second most abundant taxa in the area (26.8% broad-scale abundance), fine-scale abundance was intermediate (8.2%) and they were seldom targeted by Mediterranean lionfish (3.4%). In comparison, Labridae individuals had low broad-scale abundance (5.5%), but relatively high fine-scale abundance (15.6%) and were highly targeted (32.2%) (Table 2, Table S1).

**Table 3.**
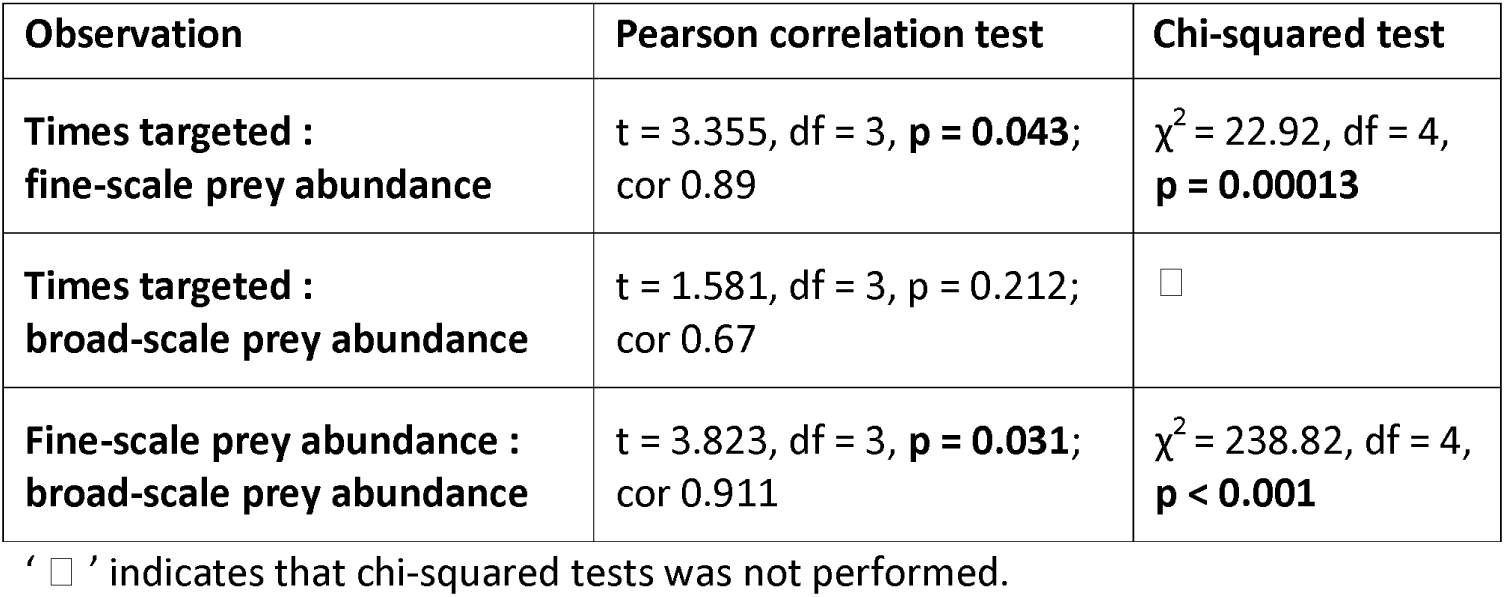
Results of Pearson correlation and chi-squared tests examining patterns of abundance of potential prey species targeted by Mediterranean lionfish, in the immediate vicinity of Mediterranean lionfish (fine-scale abundance), and in the broader area (broad-scale abundance). Correlation tests examine whether the relative abundance of prey targeted or in the vicinity of Mediterranean lionfish reflect (at least partly) the availability of those species in the community. Chi-squared tests examine whether prey species are targeted or associated with Mediterranean lionfish in proportions that are significantly different from what would be expected at random.

| Observation | Pearson correlation test | Chi-squared test |
| --- | --- | --- |
| **Times targeted :**<br>**fine-scale prey abundance** | t = 3.355, df = 3, **p = 0.043**;<br>cor 0.89 | $\chi^2 = 22.92$, df = 4,<br>**p = 0.00013** |
| **Times targeted :**<br>**broad-scale prey abundance** | t = 1.581, df = 3, p = 0.212;<br>cor 0.67 | □ |
| **Fine-scale prey abundance :**<br>**broad-scale prey abundance** | t = 3.823, df = 3, **p = 0.031**;<br>cor 0.911 | $\chi^2 = 238.82$, df = 4,<br>**p < 0.001** |

' □ ' indicates that chi-squared tests was not performed.

### 3.3. Native versus invasive prey naiveté to Mediterranean lionfish predation

Origin (i.e. native versus invasive) had a significant effect on closest approach distance, while Mediterranean lionfish size and the interaction between size and origin had no significant effect (Table 4). Native fish species approached more closely to Mediterranean lionfish than invasive fish species (Fig. 4). Particularly, *C. chromis* (n =182), *T. pavo* (n = 48), *C. julis* (n = 8), *A. imberbis* (n = 3) showed the nearest approach distances ([mean cm ± SE], 25.2 ± 0.9, 32.9 ± 1.6, 23.9 ± 4.2, 23.3 ± 2.4, respectively), while *S. rivulatus* (n = 30) and *S. cretense* (n = 6) showed the furthest approach distances (53.9 ± 0.8 and 51.3 ± 1.8, respectively). 12 instances were recorded in which native prey fishes (11 *C. chromis*, one *T. pavo*) swam within less than 10 cm of a lionfish’s mouth (minimum distance 5 cm), while the closest distance recorded by an invasive prey fish species (*P. forsskali*, Mullidae) was 30 cm. In addition, one individual of *G. vittatus* showed nearly no response to approaching Mediterranean lionfish (closest approach distance ≤ 2 cm) and was eventually cornered and predated upon (supplementary materials Video 4).

**Fig. 4.**
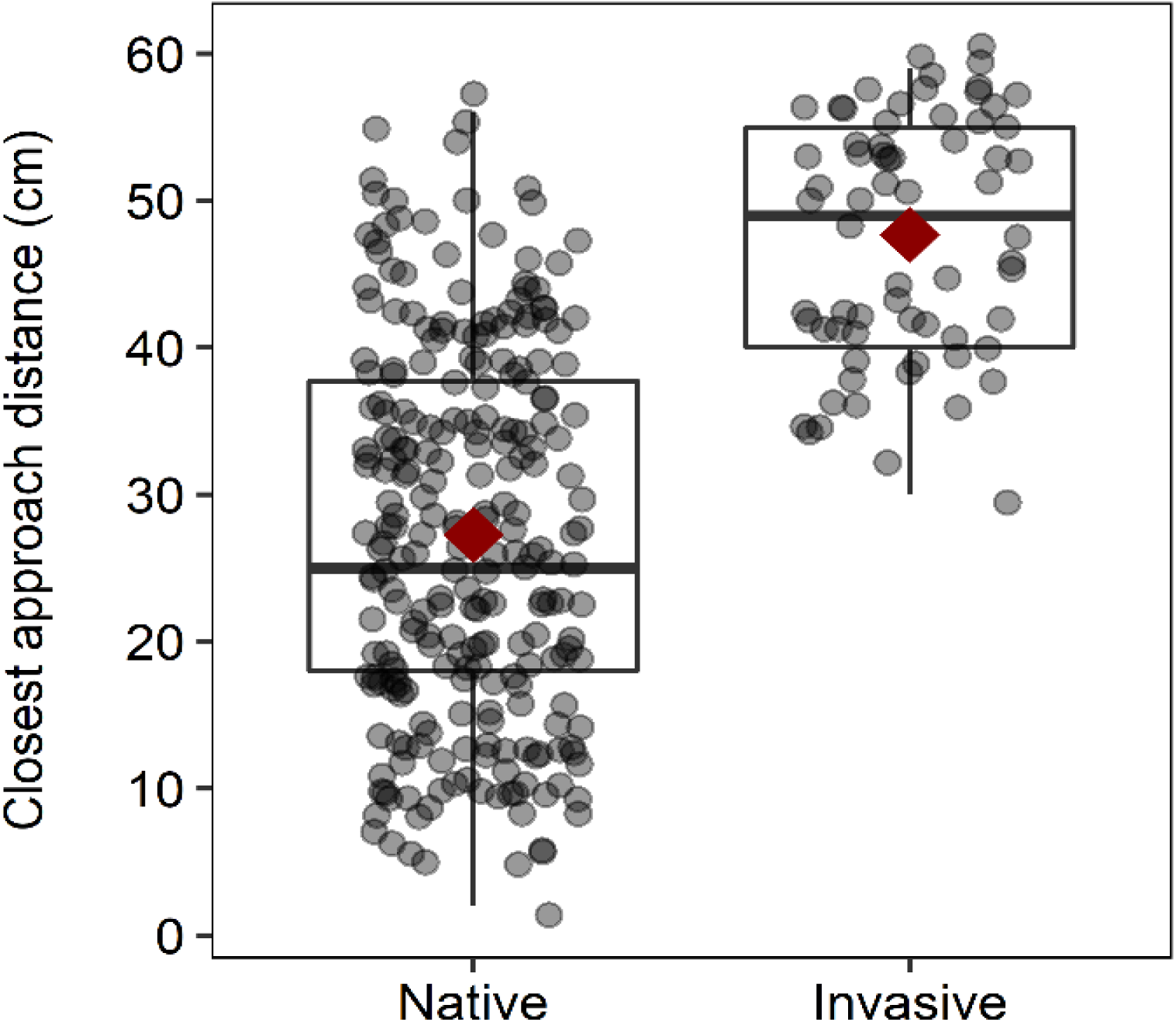
Box plot of closest approach distance (cm) of native and invasive prey fishes. See Fig. 3 for explanation of the box plots.

**Table 4.** Result of GLMM analysis, with Poisson error distribution and log link function, of the effects of origin and Mediterranean lionfish size on prey closest approach distance. Significant p values highlighted in bold.

| Response | Predictor | Residual deviance(df) | Change in deviance (df) | p |
| --- | --- | --- | --- | --- |
| **Closest approach distance** | Origin | 3171.1 (3) | 238.98 (1) | **< 0.001** |
|  | Lionfish size | 2932.1 (5) | 2.6976 (2) | 0.26 |
|  | Origin : lionfish size | 2929.4 (7) | 2.4139 (2) | 0.299 |

## 4. DISCUSSION

Given the significant impacts of lionfish invasion on native fish communities in the western Atlantic and Caribbean Sea, the recent establishment of lionfish in the Western Mediterranean has created considerable concern amongst conservationists and other sectors of the society (Kletou et al. 2016, Jimenez et al. 2017). Although the extensive array of research on the ecological, behavioural and life-history traits determining the invasive success of Atlantic lionfish may give us an indication of the consequences of the Mediterranean lionfish invasion (Côté & Smith 2018), it is important to establish whether such key traits are similarly present in the Mediterranean invasion. We examined Mediterranean lionfish in situ behaviour and feeding ecology, and also investigated whether native Mediterranean prey species are naïve towards the invasive lionfish. We found that Mediterranean lionfish, at least in shallow habitats, are crepuscular, generalist predators with some level of prey specialization. In addition, native fish species show a lower level of predator awareness to Mediterranean lionfish than invasive Lessepsian species. Our results show considerable similarities between Atlantic and Mediterranean lionfish traits and give an indication of the potential scope of impact due to the lionfish invasion, as well as the possible flow-on consequences to the ecosystem and coastal food web.

### 4.1. Mediterranean lionfish behaviour, feeding ecology and prey association

Mediterranean lionfish activity varied significantly throughout the day, though with highest activity during sunrise and sunset, with Mediterranean lionfish thus able to be described as active crepuscular predators. Such feeding behaviour is consistent with previous studies on lionfish behaviour in their native (Indo-Pacific) and invaded range (western Atlantic ocean) (Cure et al. 2012, Benkwitt 2016), suggesting that Mediterranean lionfish have maintained their native behaviour, and that activity is influenced by light levels (Côté & Maljkovic 2010). At sunrise and sunset all activities related to foraging (time spent hunting, distance moved) were highest, while around noon most of the Mediterranean lionfish were observed resting or hiding under rock ledges. In lionfish, crepuscular feeding activity may be associated with increased hunting success of lionfish due to their high visual acuity in low light conditions (Green et al. 2011). In the Mediterranean the ability of lionfish to successfully hunt during twilight hours may potentially be an additional trait leading to their invasive success. As prey abundance may increase during twilight, due to the change-over between diurnal and nocturnal species (Hobson 1973, Potts 1990), a crepuscular generalist predator will have access to a broad array of prey species, potentially increasing the likelihood of successfully colonising a new environment. Although we did not systematically quantify nocturnal activity, four active Mediterranean lionfish were observed during a pilot observation taken immediately after sunset, indicating individuals may be hunting at night. This hypothesis may be corroborated by the common findings of nocturnal prey species in Mediterranean lionfish stomach contents (Zannaki et al. 2019, Nir Stern personal communication) and by studies showing nocturnal behaviour in Red Sea lionfish (Fishelson 1975, McTee & Grubich 2014). However, studies on invasive Atlantic lionfish report little or no nocturnal activity (Green et al. 2011, McCallister et al. 2018), suggesting potential differences in nocturnal behaviour between lionfish in natal vs invaded habitats. Further studies are needed to shed light on this potentially important behavioural trait.

The likelihood of hunting behaviour was positively influenced, though not absolutely determined, by fine-scale prey abundance, with such behaviour more prevalent when prey was locally abundant. For example, high levels of predation on *C. chromis* were found, but was also expected, as this prey species was the most abundant both at the fine- and broad-scale level. The Labridae (predominantly juvenile *T. pavo*) were also heavily targeted; although the broad-scale abundance of this group was low, they showed high abundance in the vicinity of Mediterranean lionfish. Such density dependent feeding behaviour may then indicate that Mediterranean lionfish may be associating with habitats where potential prey are more locally abundant. Indeed, Hunt et al. (2019) have recently described how Atlantic lionfish’s position on the reef and aggregating behaviour are predominantly driven by large-scale habitat complexity, with such complexity resulting in high prey abundance. Such patterns highlight that lionfish are generalist predators, that may consume available and easily accessible prey (Peake et al. 2018). We can then expect that population expansion of Mediterranean lionfish to be a stepwise progression, with individuals moving with prey abundance, resulting in localised depletion of prey populations proceeded by movement of lionfish populations.

Small body size combined with a shallow body shape, and solitary, demersal behaviour may increase the likelihood of being predated on by Mediterranean lionfish. Such patterns of predation mirror those found for Atlantic lionfish (Green & Côté 2014, Chappell & Smith 2016). For example, in the current study, the heavily predated species *T. pavo* and the other Labridae juveniles fit the description of a small and shallow body shape combined with demersal behaviour, while the most heavily predated on species *C. chromis* are small sized and gather over the rocky substratum to feed on meroplankton at dusk, to shelter during the night and to breed (Harmelin 1987, Aguzzi et al. 2013, Pinnegar 2018). In comparison, despite juveniles of *S. rivulatus* and *S. cretense* being the second and third most abundant species at the broad-scale level, with intermediate fine-scale abundance, both were not an important prey species for Mediterranean lionfish. Both species are relatively deep bodied, showing schooling behaviour and (*S. rivulatus* only) defensive venomous dorsal spines (Popper & Gundermann 1975, De Girolamo et al. 1999), potentially reducing their likelihood of being preferentially preyed upon.

### 4.2. Native versus invasive prey naiveté to Mediterranean lionfish predation

Species-specific anti-predatory behaviour may influence predation rates of Mediterranean lionfish, with native Mediterranean fishes (which have been evolutionarily isolated from lionfish) showing closer approach distances (i.e. higher naiveté) to Mediterranean lionfish than co-evolved (Lessepsian) fishes. These results are in line with the native prey naiveté theory, which posits that native prey lack effective anti-predatory responses to a novel predator, due to lack of exposure to the predator archetypes over evolutionary time (Cox & Lima 2006, Sih et al. 2010, Paolucci et al. 2013). This theory has been postulated to explain the predatory advantage of lionfish in the western Atlantic Ocean compared to native predators, and the disproportionate impact that lionfish have exerted on native Atlantic prey populations (Anton et al. 2016, Haines & Côté 2019). These results suggest that, without considerable changes in the behavioural response of prey fishes in the Mediterranean, substantial negative impacts on prey populations may occur (Strauss et al. 2006). For example, a sharp decline in prey populations’ abundance will lead to reduced genetic variation, thereby diminishing the potential for adaptation to current environmental change (i.e. topicalization of Mediterranean Sea) (Strauss et al. 2006, Bianchi 2007). However, due to a strong selective predatory pressure, we would expect that prey recognition might eventually evolve in native Mediterranean fish (Strauss et al. 2006), as evolution of prey response to invasive predator has been shown to be possible in just a few generations (Berger 2001). Indeed, negative size-selective mortality (as per the highly targeted Labridae), has been linked to the selection of faster growing phenotypes (Belk et al. 1993, Ellis & Gibson 1995, Sogard 1997) coupled with the selection of shier and less active individuals (Sbragaglia et al. 2019). However, whether larger and more wary fish will be better adapted to cope with Mediterranean lionfish predation in the future is unknown. Nevertheless, to date, there is no evidence of adaptation of western Atlantic prey fish to the threat of Atlantic lionfish, even though lionfish populations have been established for over a decade (Anton et al. 2016, Haines & Côté 2019).

### 4.3. Implication for the wider marine community

As both *C. chromis* and *T. pavo* play fundamental roles in the Mediterranean ecosystem (Guidetti & Dulčić 2007, Milazzo et al. 2011, Galasso et al. 2015, Pinnegar 2018), selective predation and localised reduction in populations may have substantial flow-on effects on the wider marine community. For example, *C. chromis* populations channel carbon, phosphorus and nitrogen from the pelagic-zooplankton system to the rocky littoral environment in the form of liquid and solid waste (Bracciali et al. 2012, Pinnegar 2018), subsidising nutrients in one of the most oligotrophic seas in the world (Krom et al. 2014).

Furthermore, *C. chromis* is the most abundant prey species of meso-predator fish and seabirds, and a major consumer of zooplankton and fish eggs itself (Pinnegar 2018), meaning that reductions in the abundance of *C. chromis* may have a strong negative impact on the coastal food web. In parallel, *T. pavo* feeds on small molluscs, crustaceans, annelids and echinoderms (Guidetti 2004, Galasso et al. 2015, Sinopoli et al. 2017), with its population abundance having been linked to top-down regulation of sea urchin populations (Guidetti & Dulčić 2007, Galasso et al. 2015). Therefore, reductions in the abundance of this species may accelerate local shifts in community structure from the ubiquitous sea grass bed to predominantly grazed barrens, in an environment already under strong pressure by invasive fish grazers (i.e. *Siganus rivulatus* and *S. luridus*) (Vergés et al. 2014). Lastly, Mediterranean lionfish have been increasingly shown to utilise caves as diurnal refugee habitats in Cyprus and, perhaps, feeding grounds (Jimenez et al. 2019b). Such behaviour may then overlap and, through increased predation, impact the diel migrations of fish and crustacean species between caves and external habitats. Such migration provide an important source of nutrients to the relatively nutrient poor epibenthic communities in caves (Coma et al. 1997, Bussotti et al. 2018). The impact of Mediterranean lionfish preying on diel migrants may then have substantial consequences for the maintenance of such nutrient poor communities, increasing the likelihood of locally mediated losses of cave specific benthic communities.

Our findings show that Mediterranean lionfish possess many of the behavioural and ecological traits that have been key in determining the invasive success of the Atlantic lionfish. Although the Mediterranean lionfish invasion is still at its early stage (first reported established populations in Lebanon and Cyprus in 2015) (Jimenez et al. 2016, Azzurro & Bariche 2017), selective predation and prey naiveté may have substantial population impacts on a range of native prey species (Albins & Hixon 2013, Peake et al. 2018, Côté & Smith 2018). Such impacts are expected to be in line with biodiversity impacts of the western Atlantic lionfish invasion, which has been linked to significant changes in native fish community, including reductions in prey species density, biomass, recruitment, richness (Albins & Hixon 2008, Benkwitt 2015, Albins 2015, Palmer et al. 2016), and even local species extirpation (Ingeman 2016). Ultimately, as the Mediterranean lionfish are at the beginning of the spreading stage of the invasion process (Chapple et al. 2012), is still unknown whether the behavioural and life-history traits measured in this study will be adaptive for a pan-Mediterranean invasion, where different thermal-regime and oceanographic conditions occur (Johnston & Purkis 2014). Hence, further studies are needed to track the potential expansion of Mediterranean lionfish distribution, and to evaluate the potential for predator-prey co-evolution of life-history and behavioural traits, by comparing differences in those traits between well-established and recently invaded areas.

## ACKNOWLEDGEMENTS

We thank the University of Nottingham for the support throughout the whole study. Marios Josephides and the Department of Fisheries and Marine Research (DFMR) of Cyprus for funding the underwater fish surveys and providing data, and Pantelis Patsalou for fieldwork and logistic support. We also thank ADC MacColl and the anonymous reviewers for precious advices and constructive suggestions. This work was funded by D. D’Agostino’s PhD scholarship (Nottingham University), Enalia Physis and the operational program ‘Thalassa 2014-2020’ (75% by the European Maritime and Fisheries Fund & 25% by national funds).

## LITERATURE CITED

Aguzzi J, Sbragaglia V, Santamaría G, Del Río J, Sardà F, Nogueras M, Manuel A (2013) Daily activity rhythms in temperate coastal fishes: insights from cabled observatory video monitoring. Mar Ecol Prog Ser 486:223–236.

Albins M (2015) Invasive Pacific lionfish Pterois volitans reduce abundance and species richness of native Bahamian coral-reef fishes. Mar Ecol Prog Ser 522:231–243.

Albins MA, Hixon MA (2008) Invasive Indo-Pacific lionfish Pterois volitans reduce recruitment of Atlantic coral-reef fishes. Mar Ecol Prog Ser 367:233–238.

Albins MA, Hixon MA (2013) Worst case scenario: Potential long-term effects of invasive predatory lionfish (Pterois volitans) on Atlantic and Caribbean coral-reef communities. Environ Biol Fishes 96:1151–1157.

Anton A, Cure K, Layman CA, Puntila R, Simpson MS, Bruno JF (2016) Prey naiveté to invasive lionfish Pterois volitans on Caribbean coral reefs. Mar Ecol Prog Ser 544:257–269.

Azzurro E, Bariche M (2017) Local knowledge and awareness on the incipient lionfish invasion in the eastern Mediterranean Sea. Mar Freshw Res 68:1950.

Azzurro E, Stancanelli B, Di Martino V, Bariche M (2017) Range expansion of the common lionfish Pterois miles (Bennett, 1828) in the Mediterranean Sea: an unwanted new guest for Italian waters. BioInvasions Rec 6:95–98.

Azzurro E, Tuset VM, Lombarte A, Maynou F, Simberloff D, Rodríguez-Pérez A, Solé R V. (2014) External morphology explains the success of biological invasions. Ecol Lett 17:1455–1463.

Bariche M; Torres, M.; Azzurro E (2013) The presence of the invasive Lionfish Pterois miles in the Mediterranean Sea. Mediterr Mar Sci 14:292–294.

Belk MC, Hales LS, Belk MC, Hales LS (1993) Predation-Induced Differences in Growth and Reproduction of Bluegills (Lepomis macrochirus). Copeia:1034–1044.

Benkwitt CE (2016) Invasive lionfish increase activity and foraging movements at greater local densities. Mar Ecol Prog Ser 558:255–266.

Benkwitt CE (2015) Non-linear effects of invasive lionfish density on native coral-reef fish communities. Biol Invasions 17:1383–1395.

Benkwitt CE (2017) Predator effects on reef fish settlement depend on predator origin and recruit density. Ecology 98:896–902.

Berger J (2001) Recolonizing Carnivores and Naive Prey: Conservation Lessons from Pleistocene Extinctions. Science (80) 291:1036–1039.

Bianchi CN (2007) Biodiversity issues for the forthcoming tropical Mediterranean Sea. Hydrobiologia 580:7–21.

Bracciali C, Campobello D, Giacoma C, Sarà G (2012) Effects of nautical traffic and noise on foraging patterns of mediterranean Damselfish (Chromis chromis). PLoS One 7.

Bussotti S, Di Franco A, Bianchi CN, Chevaldonné P, Egea L, Fanelli E, Lejeusne C, Musco L, Navarro-Barranco C, Pey A, Planes S, Vieux-Ingrassia JV, Guidetti P (2018) Fish mitigate trophic depletion in marine cave ecosystems. Sci Rep 8:9193.

Chappell BF, Smith KG (2016) Patterns of predation of native reef fish by invasive Indo-Pacific lionfish in the western Atlantic: Evidence of selectivity by a generalist predator. Glob Ecol Conserv 8:18–23.

Chapple DG, Simmonds SM, Wong BBM (2012) Can behavioral and personality traits influence the success of unintentional species introductions? Trends Ecol Evol 27:57–64.

Coma R, Carola M, Riera T, Zabala M (1997) Horizontal Transfer of Matter by a Cave-Dwelling Mysid. Mar Ecol 18:211–226.

Côté IM, Maljkovic A (2010) Predation rates of Indo-Pacific lionfish on Bahamian coral reefs. Mar Ecol Prog Ser 404:219–225.

Côté IM, Smith NS (2018) The lionfish Pterois sp. invasion: Has the worst-case scenario come to pass? J Fish Biol 92:660–689.

Cox JG, Lima SL (2006) Naiveté and an aquatic-terrestrial dichotomy in the effects of introduced predators. Trends Ecol Evol 21:674–680.

Crutzen PJ (2006) The anthropocene. In: Ehlers E, Krafft T (Eds) Earth System Science in the Anthropocene. Springer Berlin Heidelberg, p 13–18

Cure K, Benkwitt C, Kindinger T, Pickering E, Pusack T, McIlwain J, Hixon M (2012) Comparative behavior of red lionfish Pterois volitans on native Pacific versus invaded Atlantic coral reefs. Mar Ecol Prog Ser 467:181–192.

D’Agostino D, Burt JA, Reader T, Vaughan GO, Chapman BB, Santinelli V, Cavalcante GH, Feary DA (2019) The influence of thermal extremes on coral reef fish behaviour in the Arabian/Persian Gulf. Coral Reefs doi:10.1007/s00338-019-01847-z.

Dirzo R, Young HS, Galetti M, Ceballos G, Isaac NJB, Collen B (2014) Defaunation in the Anthropocene. Science 345:401–6.

Edwards MA, Frazer TK, Jacoby CA (2014) Age and growth of invasive lionfish (Pterois spp.) in the Caribbean Sea, with implications for management. Bull Mar Sci 90:953–966.

Ellis T, Gibson RN (1995) Size-selective predation of 0-group flatfishes on a Scottish coastal nursery ground. Mar Ecol Prog Ser 127:27–37.

Felden A, Paris CI, Chapple DG, Haywood J, Suarez A V., Tsutsui ND, Lester PJ, Gruber MAM (2018) Behavioural variation and plasticity along an invasive ant introduction pathway. J Anim Ecol 87:1653–1666.

Galanidi M, Zenetos A, Bacher S (2018) Assessing the socio-economic impacts of priority marine invasive fishes in the Mediterranean with the newly proposed SEICAT methodology. Mediterr Mar Sci 19:107.

Galasso NM, Bonaviri C, Trapani F Di, Picciotto M, Gianguzza P, Agnetta D, Badalamenti F (2015) Fish-seastar facilitation leads to algal forest restoration on protected rocky reefs. Sci Rep 5:1–9.

Gallardo B, Clavero M, Sánchez MI, Vilà M (2016) Global ecological impacts of invasive species in aquatic ecosystems. Glob Chang Biol 22:151–163.

García-Rivas MC, Machkour-M’Rabet S, Pérez-Lachaud G, Schmitter-Soto JJ, Cé réghino R, Doneys C, St-Jean N, Hénaut Y (2018) Age-dependent strategies related to lionfish activities in the Mexican Caribbean. Environ Biol Fishes 101:563–578.

de Girolamo M, Scaggiante M, Rasotto MB (1999) Social organization and sexual pattern in the Mediterranean parrotfish Sparisoma cretense (Teleostei: Scaridae). Mar Biol 135:353–360.

Gordon TAC, Harding HR, Clever FK, Davidson IK, Davison W, Montgomery DW, Weatherhead RC, Windsor FM, Armstrong JD, Bardonnet A, Bergman E, Britton JR, Côté IM, D’Agostino D, Greenberg LA, Harborne AR, Kahilainen KK, Metcalfe NB, Mills SC, Milner NJ, Mittermayer FH, Montorio L, Nedelec SL, Prokkola JM, Rutterford LA, Salvanes AGV, Simpson SD, Vainikka A, Pinnegar JK, Santos EM (2018) Fishes in a changing world: learning from the past to promote sustainability of fish populations. J Fish Biol 92:804–827.

Green S, Akins J, Côté I (2011) Foraging behaviour and prey consumption in the Indo-Pacific lionfish on Bahamian coral reefs. Mar Ecol Prog Ser 433:159–167.

Guidetti P (2004) Consumers of sea urchins, Paracentrotus lividus and Arbacia lixula, in shallow Mediterranean rocky reefs. Helgol Mar Res 58:110–116.

Guidetti P, Dulčić J (2007) Relationships among predatory fish, sea urchins and barrens in Mediterranean rocky reefs across a latitudinal gradient. Mar Environ Res 63:168–184.

Haines LJ, Côté IM (2019) Homing decisions reveal lack of risk perception by Caribbean damselfish of invasive lionfish. Biol Invasions 21:1657–1668.

Hayes KR, Barry SC (2008) Are there any consistent predictors of invasion success? Biol Invasions 10:483–506.

Hobson ES (1973) Diel feeding migrations in tropical reef fishes. Helgol∼inder wiss Meeresunters 24:361–370.

Holway DA, Suarez A V (1999) Animal behavior!]: an essential component of invasion biology. Trends Ecol Evol 14:328–330.

Hunt CL, Kelly GR, Windmill H, Curtis-Quick J, Conlon H, Bodmer MDV, Rogers AD, Exton DA (2019) Aggregating behaviour in invasive Caribbean lionfish is driven by habitat complexity. Sci Rep 9:1–9.

Ingeman KE (2016) Lionfish cause increased mortality rates and drive local extirpation of native prey. Mar Ecol Prog Ser 558:235–245.

Jimenez C, Andreou V, Hadjioannou L, Petrou A, Alhaija RA, Patsalou P (20 17) Not everyone’s cup of tea: Public perception of culling invasive lionfish in Cyprus. J Black Sea/Mediterranean Environ 23:38–47.

Jimenez C, Petrou A, Andreou V, Hadjioannou L, Wolf W, Koutsoloukas N, Alhaija RA (2016) Veni, vidi, vici: the successful establishment of the lionfish pterois miles in Cyprus (Levantine Sea). Rapid Commun int Mer Méditerranea 41:417.

Johnston MW, Purkis SJ (2014) Are lionfish set for a Mediterranean invasion? Modelling explains why this is unlikely to occur. Mar Pollut Bull 88:138–147.

Kleitou P, Hall-Spencer J, Rees S, Sfenthourakis S, Demetriou A, Chartosia N, Jimenez C, Hadjioannou L, Petrou A, Christodoulides Y, Georgiou A, Andreou V, Antoniouc C, Savva I, Kletou D (2019) Tackling the lionfish invasion in the Mediterranean. The EU-LIFE RELIONMED Project!]: progress and results of the 1st Mediterranean Symposium on the Non-Indigenous Species, Antalya, Turkey, 17-19 January 2019. proceeding msnis.

Kletou D, Hall-Spencer JM, Kleitou P (2016) A lionfish (Pterois miles) invasion has begun in the Mediterranean Sea. Mar Biodivers Rec 9:46.

Kolar CS, Lodge DM (2001) Progress in invasion biology: predicting invaders. Trends Ecol Evol 16:199–204.

McCallister M, Renchen J, Binder B, Acosta A (2018) Diel Activity Patterns and Movement of Invasive Lionfish (Pterois volitans/P. miles) in the Florida Keys Identified Using Acoustic Telemetry. Gulf Caribb Res 29:27–40.

McGill BJ, Dornelas M, Gotelli NJ, Magurran AE (2015) Fifteen forms of biodiversity trend in the anthropocene. Trends Ecol Evol 30:104–113.

McTee SA, Grubich JR (2014) Native densities, distribution, and diurnal activity of Red Sea lionfishes (Scorpaenidae). Mar Ecol Prog Ser 508:223–232.

Milazzo M, Palmeri A, Falcón JM, Badalamenti F, Garcìa-Charton JA, Sinopoli M, Chemello R, Brito A (2011) Vertical distribution of two sympatric labrid fishes in the Western Mediterranean and Eastern Atlantic rocky subtidal: local shore topography does matter. Mar Ecol 32:521–531.

Morris JA, Whitfield PE (2009) Biology, ecology, control and management of the invasive Indo-Pacific lionfish: an updated integrated assessment. NOAA Technical Memorandum NOS NCCOS 99.

Moyle PB, Light T (1996) Biological invasion of fresh water: empirical rules and assembly theory. Biol Conserv 78:149–161.

Palmer G, Hogan JD, Sterba-Boatwright BD, Overath RD (2016) Invasive lionfish pterois volitans reduce the density but not the genetic diversity of a native reef fish. Mar Ecol Prog Ser 558:223–234.

Paolucci EM, MacIsaac HJ, Ricciardi A (2013) Origin matters: alien consumers inflict greater damage on prey populations than do native consumers. Divers Distrib 19:988–995.

Peake J, Bogdanoff AK, Layman CA, Castillo B, Reale-Munroe K, Chapman J, D ahl K, Patterson III WF, Eddy C, Ellis RD, Faletti M, Higgs N, Johnston MA, Muñoz RC, Sandel V, Villasenor-Derbez JC, Morris JA (2018) Feeding ecology of invasive lionfish (Pterois volitans and Pterois miles) in the temperate and tropical western Atlantic. Biol Invasions 20:2567–2597.

Pinnegar JK (2018) Why the damselfish Chromis chromis is a key species in the Mediterranean rocky littoral – a quantitative perspective. J Fish Biol 92:851–872.

Popper D, Gundermann N (1975) Some ecological and behavioural aspects of siganid populations in the Red Sea and Mediterranean coasts of Israel in relation to their suitability for aquaculture. Aquaculture 6:127–141.

Potts GW (1990) Crepuscular behaviour of marine fishes. In: Herring PJ, Campbell AK, Whitfield M, Maddock L (Eds). Light and life in the sea. Cambridge University Press, Cambridge, 221–227.

Sbragaglia V, Al J, Fromm K, Monk CT, Uusi-heikkil S, Honsey AE, Wilson ADM (2019) Experimental Size-Selective Harvesting Affects Behavioral Types of a Social Fish os. Trans Am Fish Soc 148:552–568.

Sih A, Bolnick DI, Luttbeg B, Orrock JL, Peacor SD, Pintor LM, Preisser E, Rehage JS, Vonesh JR (2010) Predator-prey naïveté, antipredator behavior, and the ecology of predator invasions. Oikos 119:610–621.

Sinopoli M, Chemello R, Vaccaro A, Milazzo M (2017) Food resource partitioning between two sympatric temperate wrasses. Mar Freshw Res 68:2324.

Sogard SM (1997) Size-selective mortality in the juvenile stage of teleost fishes: a review. Bull Mar Sci 60:1129–1157.

Strauss SY, Lau JA, Carroll SP (2006) Evolutionary responses of natives to introduced species: what do introductions tell us about natural communities? Ecol Lett 9:357–374.

Strayer DL (2010) Alien species in fresh waters!]: ecological effects, interactions with other stressors, and prospects for the future. Freshw Biol 55:152–174.

Vergés A, Steinberg PD, Hay ME, Poore AGB, Campbell AH, Ballesteros E, Heck KL, Booth DJ, Coleman MA, Feary DA, Figueira W, Langlois T, Marzinelli EM, Mizerek T, Mumby PJ, Nakamura Y, Roughan M, van Sebille E, Gupta A Sen, Smale DA, Tomas F, Wernberg T, Wilson SK (2014) The tropicalization of temperate marine ecosystems: Climate-mediated changes in herbivory and community phase shifts. Proc R Soc B Biol Sci 281: 20140846.

Zannaki K, Corsini-Foka M, Kampouris TE, Batjakas IE (2019) First results on the diet of the invasive Pterois miles (Actinopterygii: Scorpaeniformes: Scorpaenidae) in the Hellenic waters. Acta Ichthyol Piscat 49:311–317.

